# Establishment of a Chikungunya virus pseudotype system strictly dependent on viral protein expression

**DOI:** 10.1101/2024.12.19.629062

**Authors:** Atsushi Tanaka, Takayuki Miyazawa

**Affiliations:** Division of Research Animal Laboratory and Translational Medicine, Research and Development Center Osaka Medical and Pharmaceutical University, Takatsuki, Osaka 569-8686, Japan; Laboratory of Virus-Host Coevolution, Institute for Life and Medical Sciences, Kyoto University, Sakyo-ku, Kyoto 606-8507, Japan; Kyoto Animal Human Organism Research Institute, Nakagyo-ku, Kyoto, Japan

**Author notes:** Correspondence: Atsushi Tanaka.

## Abstract

Chikungunya virus (CHIKV) is an enveloped RNA virus that causes Chikungunya fever in humans. It is classified into the arboviruses (arthropod-borne viruses) and is transmitted by mosquitoes. Therefore, mosquitoes can replicate many types of cells derived from mammals or insects. In this study, we tried to establish the widely useable Chikungunya virus pseudotype-system adapting various viral species, and we demonstrated the production of Chikungunya pseudotype virus baring the envelope protein from two different viral families, Coronaviridae or Rhabdoviridae i.e., severe acute respiratory syndrome coronavirus 2 spike protein (CoV-2-S) or vesicular stomatitis virus glycoprotein (VSV-G), respectively. We found that the capsid protein of Chikungunya virus is not always necessary in the formation of Chikungunya virus-based pseudotypes, but that the capsid protein increases the efficiency of expression of the sub-genomic RNA which codes the labeled genes. Our established pseudotype virus-producing system supplied a sufficient titer of virions for application to most virological experiments that showed more than 10^4^ focus forming units (FFU)/ml. The pseudotype infections were strictly dependent on compatibility between the viral envelope protein and its receptor and there was no false-positive background infection. Our established pseudotype virus system can be used as a robust platform to study various virus infections and for screening and in-depth evaluation of neutralizing antibodies and antiviral agents.

## Introduction

Chikungunya virus (CHIKV) is a mosquito-borne alphavirus in the family Togaviridae and is the causative agent of chikungunya fever, which occurs mainly in tropical regions such as those in Africa, South Asia, and Southeast Asia [1]. Chikungunya fever usually begins 2 to 12 days after the mosquito bite. It is characterized by the sudden onset of a high fever that is frequently accompanied by severe joint pain, muscle pain, headache, nausea, fatigue, and rash, lasting for several days, and this acute phase is often followed by a chronic phase, characterized by persistent and crippling arthralgia [2-4]. During CHIKV infection, viremia lasts 5–7 days, and high viral loads of up to 10^9^ viral RNA copies per ml of blood plasma can be detected in the early stages of infection [5, 6].

CHIKV has a single positive-stranded RNA genome of 11.8 kbp encoding four nonstructural and five structural proteins. The structural proteins are translated from a subgenomic RNA as a single polyprotein, which is processed co-translationally into five structural proteins: capsid, E3, E2, 6 K, and E1 [7]. These structural proteins form two T = 4 quasi-icosahedral symmetry layers: the viral surface lipid membrane with a dia. of 65–70 nm containing 80 viral envelope spikes that consist of 240 copies of the E1-E2 heterodimer, and the icosahedral nucleocapsid core comprised of 240 copies of the capsid [8-11]. The viral envelope E2 glycoprotein is responsible for the binding to the cell surface receptor, and the E1 protein serves as a fusion protein [9, 12-17].

During assembly of CHIKV virus particles, the CHIKV genomic RNA is selectively packaged into the capsid core through interaction between a specific packaging signal in the nsP2 region of the viral genomic RNA and the capsid core[18]. The viral capsid core and envelope proteins are arranged in organized lattices linked via the interaction of the E2 cytoplasmic tail/endodomain with the capsid protein [19]. Such specific interactions between viral components allow for the effective formation of viral particles [19, 20]. The icosahedral nucleocapsid core of the alphavirus is assembled in not only the infected cells in the absence of envelope protein expression [21, 22] but also in vitro when incubated with RNA [23]. It has been reported that the lattice formed of viral envelope glycoprotein spikes can itself promote nucleocapsid formation[24-27] and the cytoplasmic nucleocapsids undergo rearrangements during budding [21, 24, 28], suggesting that interactions between the two lattices may influence the mature particle structure.

Although it has been reported that the interaction between the viral genomic RNA and the capsid protein, and the between the capsid protein and the envelope E2 protein are important for the formation of infectious virus particles, as described above, Zhang et al. recently constructed an infectious CHIKV with complete capsid deletion as a live attenuated vaccine candidate [29, 30] was indicating that these interactions were not necessarily required for viral particle formation.[29, 30].

CHIKV has been shown to infect various types of vertebrates as potential natural animal hosts as well as a wide range of cell lines in vitro [31, 32], and to target a wide range of organs and tissues, including the joints, skin, liver, muscle, and secondary lymphoid organs [32, 33]. The broad range of cell tropism that CHIKV can infect is likely due to the ubiquitous expression of putative receptors to which CHIKV envelope glycoproteins bind across a range of species and cell types [34-36], as well as a CHIKV replication system that can function efficiently across a range of species and cell types [37].

Chikungunya pseudotyped viruses have been constructed based on a lentiviral vector [38], murine retrovirus vector and recombinant vesicular stomatitis virus (VSV)-based chimeric viruses [39-41], that constructed using each viral genome for expressing labeled gene and bearing CHIKV surface envelope (E3, E2 and E1) protein. However, a CHIKV-based pseudotyped virus that expresses a labeled gene from the CHIKV genome has not been constructed. In this study, we tried to construct CHIKV-based pseudotyped viruses to establish a pseudotype virus system that produces virus in high titer, can be used in a wide range of target cells, and can be prepared conveniently. Our constructed pseudotyped virus bears an envelope protein that is from a viral family distinct from alphaviruses such as the severe acute respiratory syndrome coronavirus 2 spike protein (CoV-2-S) or vesicular stomatitis virus glycoprotein (VSV-G), and its genome has the labeled gene, which is substituted into the structural protein coding region and expressed as subgenomic RNA. This pseudotype system has the potential to serve as a robust platform for the study of various virus infections and for screening the correspondent neutralizing antibodies and antiviral agents.

## MATERIALS AND METHODS

### Cells

A baby hamster kidney fibroblast cell line (BHK) and its derivative (BHK/hACE2), an African green monkey kidney cell line (Vero), a human embryonic kidney cell line (293T) and a human lung cancer cell line (Calu-3) were maintained in Dulbecco’s modified minimum essential medium (DMEM) supplemented with 10% FBS, 50 units/ml penicillin, and 50 μg/ml streptomycin. To generate BHK cells stably expressing human angiotensin-converting enzyme 2 (hACE2; GenBank accession no. NM_001371415), which we designated BHK/hACE2, an hACE2 ORF sequence was amplified by reverse transcription-PCR from 293T first-strand cDNA synthesized using a Verso cDNA Synthesis Kit (Thermo Scientific, Rockford, IL) with a random hexamer. The sequences of the PCR primers for the hACE2 ORF were hACE2/sense (5’-ttt<u>ctcgag</u>acg**ATG**TCAAGCTCTTCCTGGCTCCTTC -3’) and hACE2/antisense (5’-aaa<u>gcggccg</u>**<u>C</u>TA**AAAGGAGGTCTGAACATCATC -3’). Their respective restriction enzyme sites, *Xho* I and *Not* I, are underlined, and the position of initiation and termination codons are shown in bold font. The hACE2 PCR products were cloned into the retroviral expression vector plasmid pLPCX retroviral vectors containing a gene encoding resistance to puromycin. VSV-G-pseudotyped murine retrovirus carrying the human ACE2 gene was generated by co-transfecting 293T cells with the pLPCX-hACE2, pMD-gag-pol [42], and a vesicular stomatitis virus glycoprotein (VSV-G)-expressing plasmid (Clontech, Palo Alto, CA), pVSV-G [43]. A murine retrovirus packaging vector, pLPCX-hACE2 (VSV-G), was used to transduce BHK-21 cells in the presence of 4 μg/mL polybrene. The transduced BHK-21 cells were selected by culture in a medium containing puromycin at 1 μg/ml for over two weeks. The hACE2 expression of BHK/hACE2 cells was confirmed by immunostaining using a VECTASTAIN ABC kit (Vector Laboratories, Burlingame, CA) with anti-ACE2 rabbit polyclonal antibody (21115-1-AP) (Proteintech Group, Rosemont, IL), as shown in Supplemental figure 1.

### Plasmids and antibodies

Genomic RNA of the CHIKV Ross strain was extracted by using a QIAamp viral RNA minikit (Qiagen, Hilden, Germany), and the first-strand cDNA was synthesized using a SuperScript III reverse transcriptase kit (Invitrogen Carlsbad, CA) with a random hexamer and/or a poly(dT)20NotIXbaI primer (5’-AAATCTAGAGCGGCCGCTTTTTTTTTTTTTTTTTTTT-3’) [41]. The CHIKV Ross strain genome sequence was introduced into the RNA synthesis start site of the pCXbsr retrovirus vector, where the region downstream of the CMV enhancer-MuLV promoter region and its structural protein region in which the envelope protein region (E) or capsid and envelope protein (CE) of this CHIKV genome was substituted with the construct NanoLuc luciferase (NLuc) gene-T2A sequence-CpG-free GFP::Bsr fusion gene (Invitrogen), designated NlucGFPbsr. To produce the viral RNA effectively, a hepatitis D virus (HDV) ribozyme sequence (Rbz) [44] was introduced into the end of the poly A region of the CHIKV genome. The structure of these tagged defective CHIKV genome expression plasmids, named pCHIKVΔE-NlucGFPbsr and pCHIKVΔCE-NlucGFPbsr, is shown in Fig. 1.

**Fig. 1.**
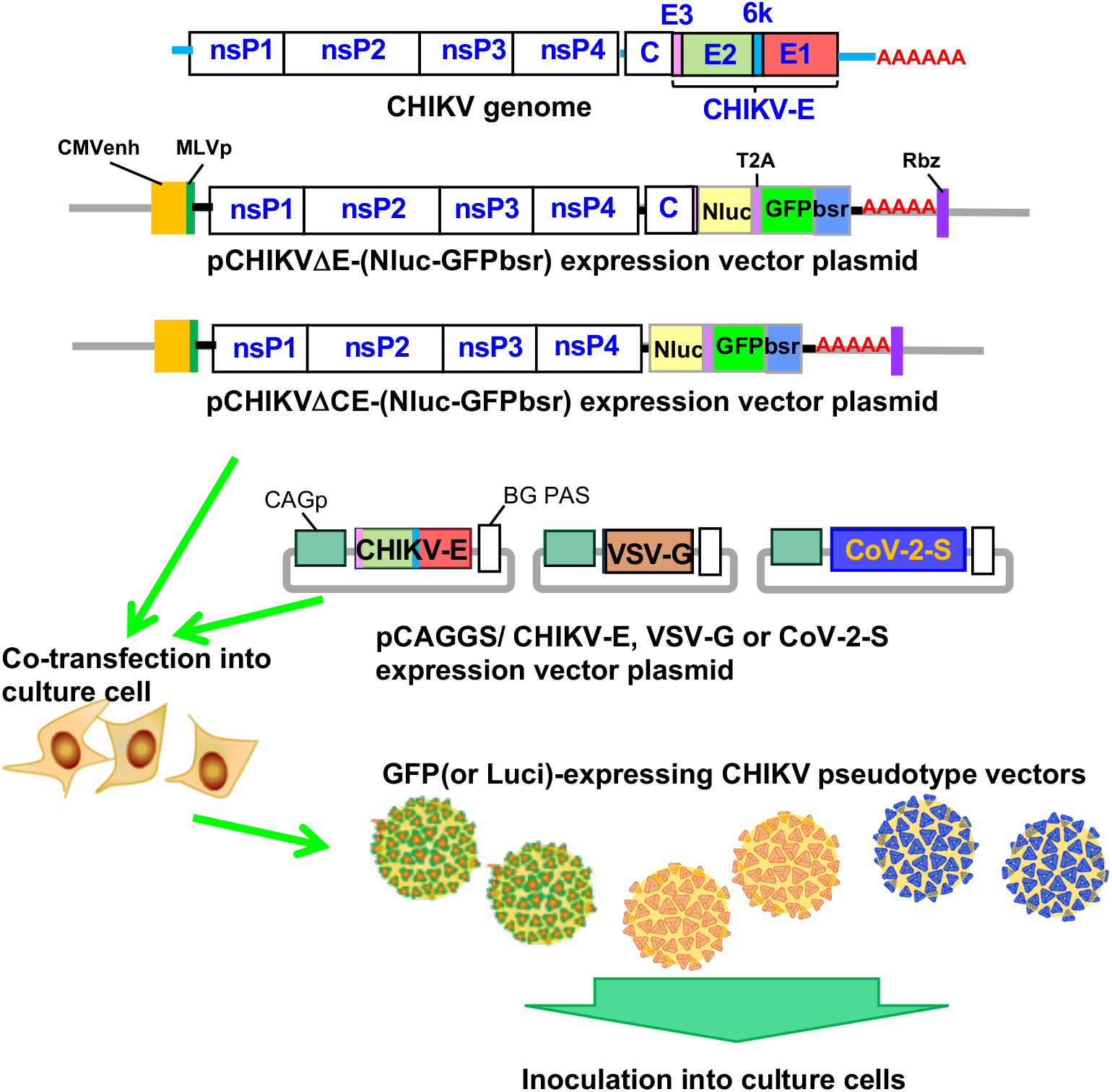
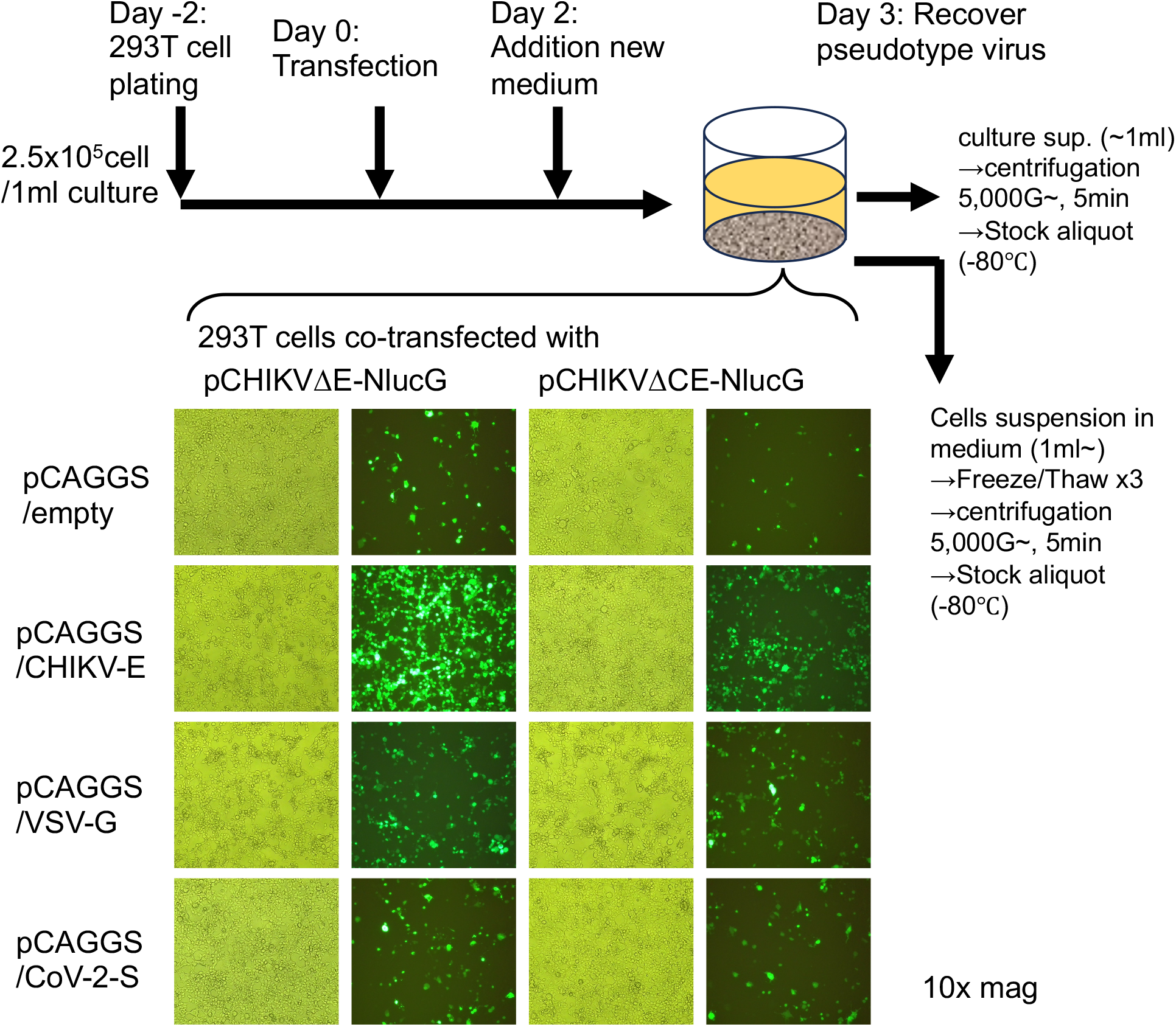
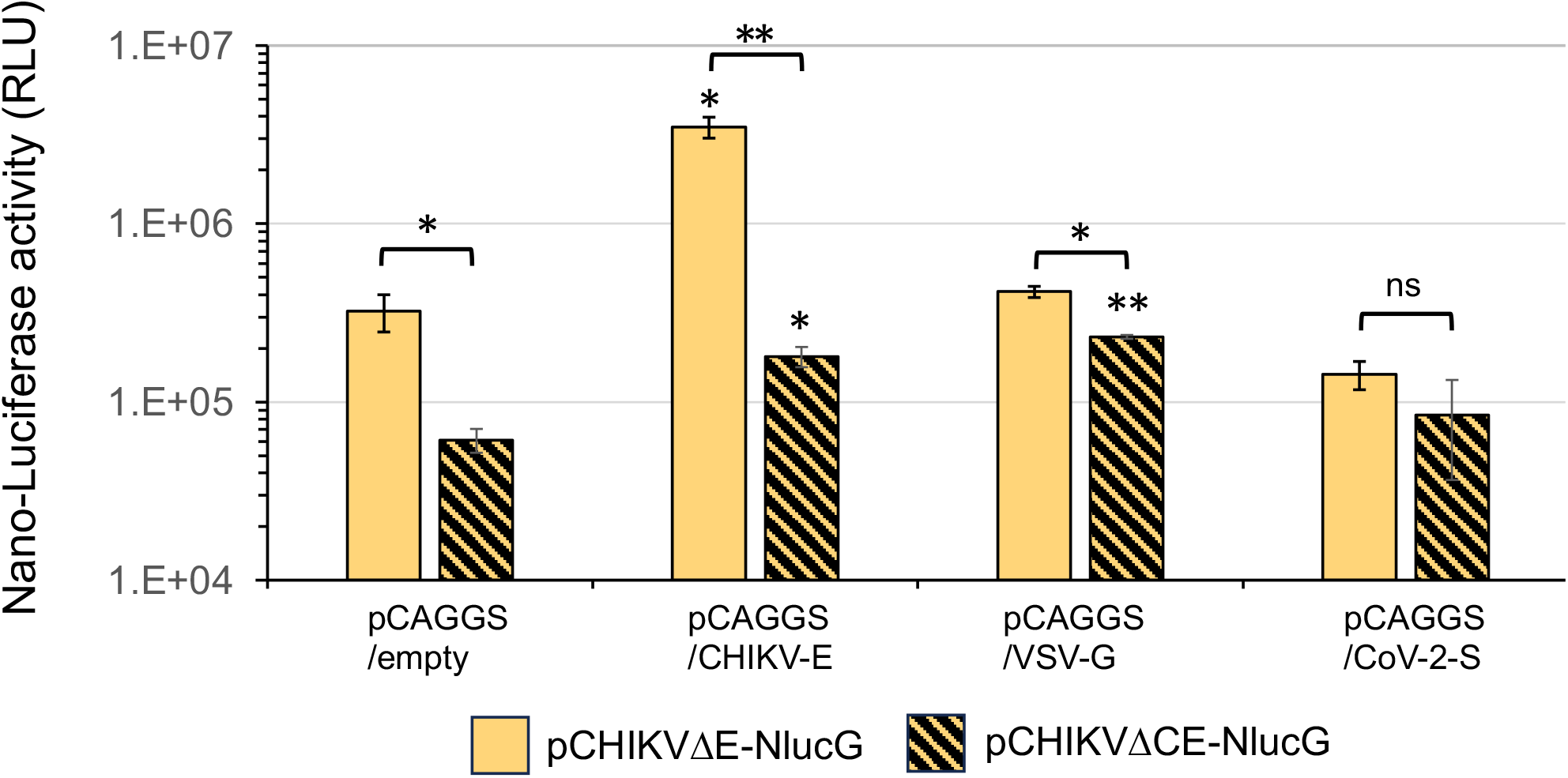
CHIKVΔE-NlucG/viral envelope protein pseudotypes design and formulation. **(A)** Shown are the tagged defective CHIKV genome expression vector plasmids, pCHIKVΔE-NlucGFPbsr and pCHIKVΔE-luc, and the viral envelope expression vectors, pCAGGS/CHIKV-E, pCAGGS/VSV-G, and pCAGGS/CoV-2-S. These expression vector plasmids were constructed as described in the Materials and Methods section. Abbreviations. nsP1-4: the nonstructural protein genes of chikungunya virus (CHIKV) strain Ross. C and E: the capsid gene (C) and envelope (E) gene (E3, E2, 6k, and E1) (i.e., the structural protein genes) of chikungunya virus (CHIKV) strain Ross. CMVenh: the human cytomegalovirus (CMV) early enhancer sequence. MLVp: the murine leukemia virus promoter sequence. Nluc: Nano luciferase gene. luc: firefly luciferase gene. T2A: thosea asigna virus 2A sequence. GFPbsr: CpG-free GFP::Bsr fusion gene (Invitrogen). HDV: hepatitis D virus. Rbz: ribozyme sequence. AAAAA: polyA sequence. CAGp: the CMV early enhancer/chicken β actin promoter of the pCAGGS expression vector. CHIKV-E, CHIKV envelope gene. VSV-G, vesicular stomatitis virus glycoprotein gene. CoV-2-S, SARS-CoV-2 spike protein gene. BG PAS, β-globin poly(A) signal. **(B)** Time course of the preparation of pseudotype samples. 293T cells were plated at 2.5x10^5^ cells/1 ml culture on 1 or 2 days before transfection (Day 0). At Day 1, 293T cells were co-transfected with pCHIKVΔE (or ΔCE)-NlucGFPbsr expression vector plasmid and pCAGGS/empty, pCAGGS/CHIKV-E, pCAGGS/VSV-G or pCAGGS/CoV-2-S vector plasmid, and then the GFP expression levels in these transfected cells were detected by fluorescence microscopy at 3 days after transfection, and then the pseudotype virus samples were harvested as the culture supernatants (Sup) and the freeze/thaw treated cells suspension with culture supernatant (F/T sup) as described in the Materials and Methods section. **(C)** Detection of NanoLuc luciferase (NLuc) activity expressed in 293T cells co-transfected with pCHIKVΔE-NlucGFPbsr and pCAGGS/ (empty, CHIKV-E, VSV-G or CoV-2-S) vector. The Nluc activity expressed in relative light units (RLU) of cell lysate equivalent to 1/500 of the volume of cell lysate in one well of a 96-well multiplate was measured, and the results shown are representative of three independent experiments and are expressed as the mean ± SD. Statistical significance was evaluated by an unpaired two-tailed *t* test. *Significant at *P* <0.05; **Significant at *P* <0.01; ns: not significant.

The CHIKV-E glycoprotein expression plasmids, pCAGGS/CHIKV-E Ross strain, were described previously [41, 45]. The VSV G protein expression plasmid, pCAGGS/VSVG, was described previously [46]. The SARS-CoV-2 wuhan strain spike protein expression plasmids, pCAGGS/CoV-2-S, encoding the codon-optimized sequence of the SARS-CoV-2 spike protein were kindly provided by Y. Matsuura (Osaka University, Osaka, Japan) [47] and the SARS-CoV-2 omicron BA.1 spike protein expression plasmids, pCAGGS/CoV-2-BA1-S, which encodes the codon-optimized sequence of the SARS-CoV-2 omicron BA.1 spike protein was synthesized by using the pCAGGS/CoV-2-S as a base and that coding the amino acid changes, deletions and insertion those different to the amino acid sequences of SARS-CoV-2 wuhan strain spike protein as follows: A67V, H69-, V70-, T95I, G142D, V143-, Y144-, Y145-, N211I, L212-, 215(insertion: EPE), G339D, R346K, S371L, S373P, S375F, K417N, N440K, G446S, S477N, T478K, E484A, Q493R, G496S, Q498R, N501Y, Y505H, T547K, D614G, H655Y, N679K, P681H, N764K, D796Y, N856K, Q954H, N969K, L981F, that showed in website: Covariant (https://covariants.org/).

The antibodies used here were as follows: anti-VSV-glycoprotein mouse monoclonal antibody, clone 8G5F11 (catalogue no. MABF2337-100UG; Merck, USA); anti-SARS-CoV-2 spike protein recombinant rabbit monoclonal antibody, clone HL257 (catalogue no. MA5-36253; ThermoFisher Scientific, Waltham, MA); anti-SARS-CoV-2 (2019-nCoV) spike neutralizing antibody, mouse monoclonal antibody, clone #43 (catalogue no. 40591-MM43; Sino Biological, Beijing, China); and anti-ACE2 rabbit polyclonal antibody (catalogue no. 21115-1-AP; Proteintech Group).

### CHIKV-based pseudotypes infection

The tagged defective CHIKV genome expression vector plasmid, pCHIKVΔE-NlucGFPbsr or pCHIKVΔCE-NlucGFPbsr, was co-transfected with the viral envelope expression vector, pCAGGS/CHIKV-E, pCAGGS/VSV-G, pCAGGS/CoV-2-S or pCAGGS/CoV-2-BA1-S, to subconfluent 293T cells that were plated 2 days before transfection. Culture supernatants and the cell suspension of cells freeze/thaw-treated 3 times with culture supernatants were harvested after 3 days of incubation at 37°C in a CO2 incubator and clarified by low-speed (∼5,000G, 5 min) centrifugation, and aliquots were stored at −80°C until use (Fig.1 B). The serially diluted 50 μl supernatants, including pseudotypes, were inoculated into the cells in the wells of a 96-well multiplate seeded with subconfluent ∼ confluent cells, and their infectivities were determined by counting GFP-positive cell foci consisting of 1–2 cells, or by measuring the NLuc activities with a Nano-Glo® Luciferase Assay System (Promega, Madison, WI) at one day after inoculation.

### CHIKV-based pseudotypes neutralization assay

The diluted samples including ∼10^2^ focus forming units (FFU) of pseudotypes were incubated with serially diluted antibodies at 37°C for 30 min, and then inoculated into the target cells in wells of a 96-well multiplate seeded with subconfluent ∼ confluent cells. After 1 day of incubation, pseudotypes titers were determined by counting GFP-positive cell foci.

## Results & Discussion

In this study, we described the establishment of a CHIKV pseudotype virus-producing system that can be applied to produce various types of envelope viruses. The indicator gene in the tagged defective CHIKV genome expression vector plasmid was expressed in not only human cells but also various types of cells from mammals and insects of different phyla [48].

In previous studies, viral samples of native CHIKV have been collected from the culture supernatant of virus-infected BHK cells, a highly productive cell line of CHIKV. CHIKV pseudotypes (murine retrovirus-based and vesicular stomatitis virus-based) bearing the CHIKV membrane envelope protein, CHIKV-E, have been collected from the culture supernatant of 293T cells transfected with pseudotype-producing gene expression plasmids [39, 41, 45]. In this study, we initially tried to prepare the pseudotypes by collecting the culture supernatants of cells co-transfected with a tagged defective CHIKV genome expression vector plasmid, pCHIKVΔE-NlucGFPbsr or pCHIKVΔCE-NlucGFPbsr, and viral envelope expression vector plasmids (Fig.1A). 293T cells transfected with pCHIKVΔE-NlucGFPbsr (or pCHIKVΔCE-NlucGFPbsr) expressed NLuc and GFP (Fig.1B, C), and the numbers of GFP-expressing cells were increased in the cells co-transfected with the CHIKV-E or VSV-G protein expression plasmid but not in those transfected with the empty or SARS-CoV-2-S protein expression plasmid, by infecting the surrounding cells with the produced pseudotypes. And also, increase of Nanoluc luciferase activity was observed in the cells co-transfected with pCHIKVΔE-NlucGFPbsr and pCAGGS/CHIKV-E; with pCHIKVΔCE-NlucGFPbsr and pCAGGS/CHIKV-E; or with pCHIKVΔCE-NlucGFPbsr and pCAGGS/VSV-G (Fig. 1C). Then, we obtained the native-form CHIKV pseudotype bearing the CHIKV-E, which we designated CHIKVΔE-NlucG/CHIKV-E, and the pseudotype bearing the VSV-G, which we named CHIKVΔE-NlucG/VSV-G, from the culture supernatants of 293T cells transfected with each, in titers of more than 10^5^ FFU/ml and more than 10^4^ FFU/ml, respectively, in inoculated BHK cells (Fig.2A).

**Fig. 2.**
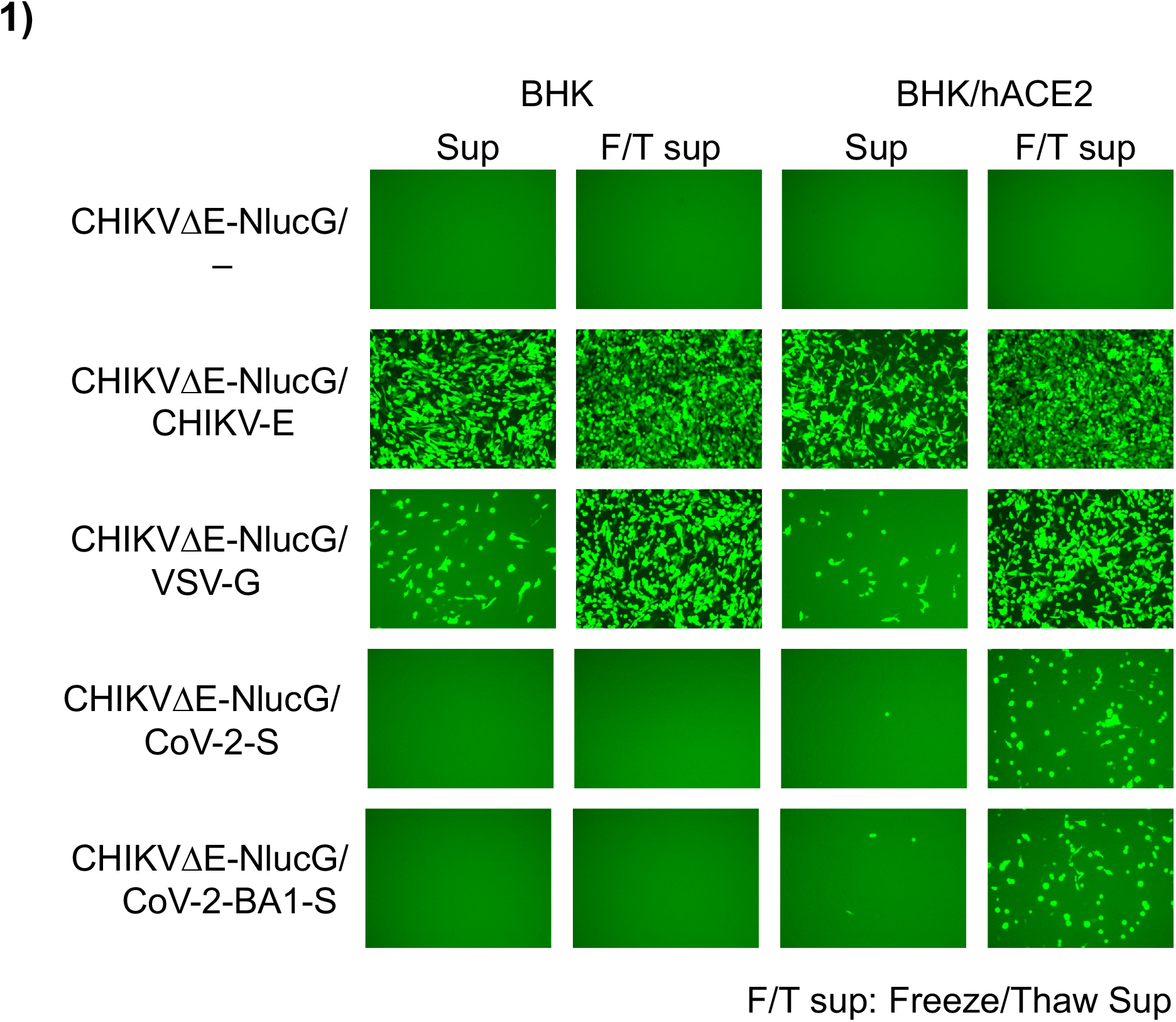
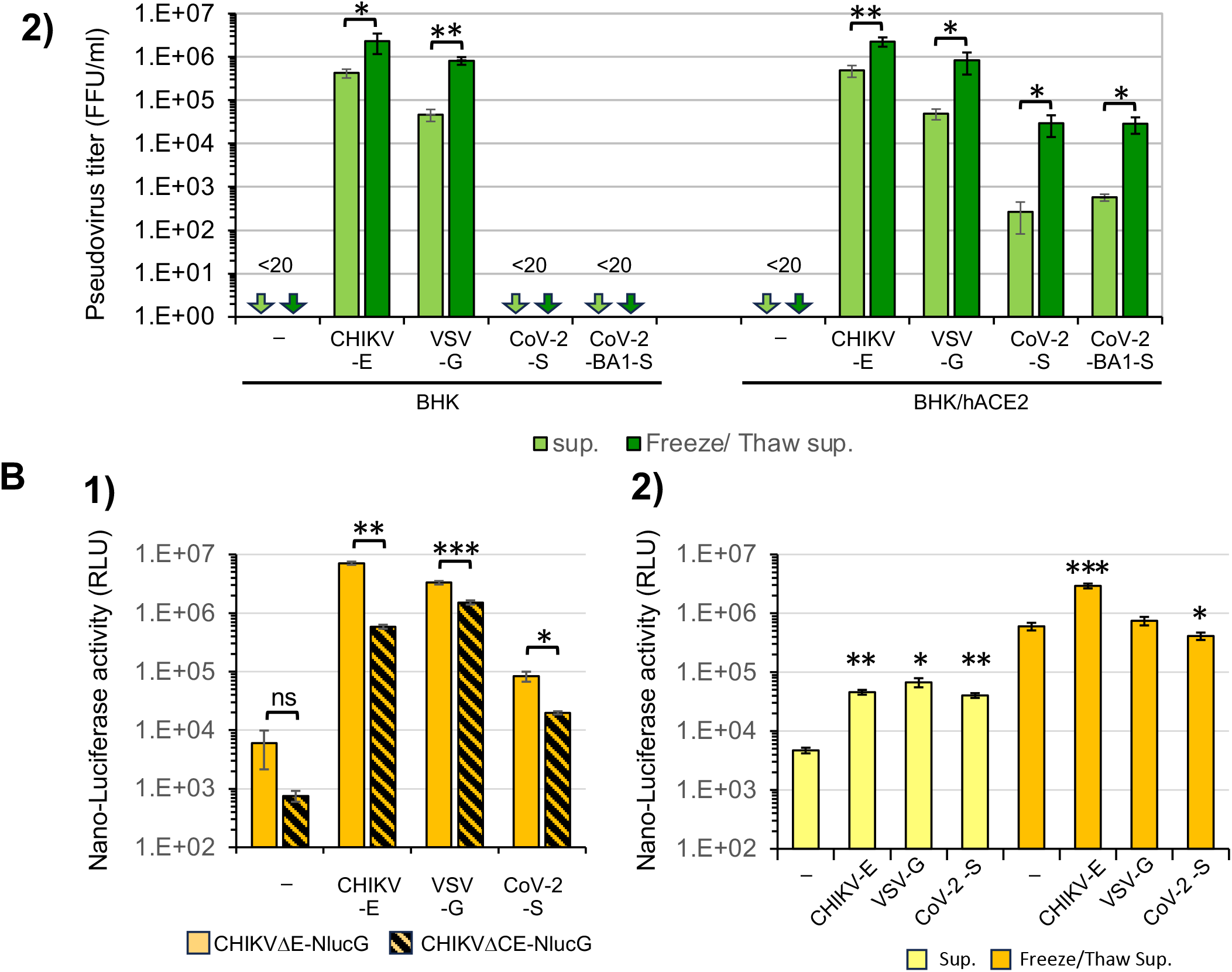
Infectivity of CHIKVΔE-NlucG/viral envelope protein pseudotypes. **(A)** CHIKVΔE-NlucG/viral envelope protein pseudotype samples obtained as Sup and F/T sup were inoculated into BHK and BHK-hACE2 cells, and GFP-expressing cells detected by fluorescence microscopy at 1 day after inoculation are shown. **(B)** (1) Detection of NLuc activity expressed in BHK/hACE2 cells infected with pCHIKVΔE (or ΔCE)-Nluc-GFPbsr/empty, CHIKV-E, VSV-G or CoV-2-S. The Nluc activity of cell lysate equivalent to 1/500 of the volume of cell lysate in one well of a 96-well multiplate was measured. (2) Detection of NLuc contained in the CHIKVΔE-NlucG/viral envelope protein pseudotype samples obtained as Sup and F/T sup. The NLuc activity of pseudotype samples were detected by using 0.2 μl equivalent of each sample. Data shown are representative of three independent experiments and are expressed as the mean ± SD. Statistical significance was evaluated by an unpaired two-tailed *t* test. *Significant at *P* <0.05; **Significant at *P* <0.01; ***Significant at *P* <0.001.

On the other hand, the BHK cells expressed no human ACE2 and were completely resistant to SARS-CoV-2 infection (Supplemental figure 2), and CHIKV pseudotype bearing the SARS-CoV-2-S proteins named CHIKVΔE-NlucG/CoV-2-S and CoV-2-BA1-S also could not infect BHK cells (Fig. 2A). Therefore, we established human ACE2-expressing BHK cells, designated BHK/hACE2 cells, that were highly susceptible to native SARS-CoV-2 (Supplemental figure 2) and then inoculated them with CHIKVΔE-NlucG/CoV-2-S (or BA1-S). Although we detected the pseudotypes bearing the SARS-CoV-2-S or BA1-S, their titers were only ∼10^2^ FFU/ml (Fig. 2A).

Previous studies reported that the viral core of alphaviruses containing the viral genomic RNA was enveloped with the cell surface plasma membrane, which contains the viral envelope protein, and budded from the cell surface [49, 50], while coronavirus genomic RNA complexed with nucleocapsid proteins was enveloped at the endoplasmic reticulum (ER)-Golgi interface and viral particles were budded into the lumen of the intermediate compartment [51-53]. Recently, Erik et al. showed that some CHIKV particles acquire an envelope intracellularly without budding from the cell surface membrane [54]. We therefore suspected that a large proportion of CHIKVΔE-NlucG/CoV-2-S accumulated at the cytoplasmic compartment. We then used freeze and thaw treatments of cells in an attempt to break the plasma membranes and release the CHIKVΔE-NlucG/CoV-2-S pseudotype virion from the cytoplasmic compartments. As expected, we were able to recover the infectious CHIKVΔE-NlucG/CoV-2-S pseudotypes from the culture supernatant of cells with plasma membranes fractured by freeze and thaw treatments at levels 100 times higher than the corresponding levels from the culture supernatant of living cells (Fig. 2). This freeze and thaw treatment of cells increased the recovery of the titers of CHIKVΔE-NlucG/CHIKV-E and CHIKVΔE-NlucG/VSV-G pseudotypes by 5- and 17-fold compared to the titers in the culture supernatants of living cells. The pseudotypes bearing no viral envelope protein, CHIKVΔE-NlucG/- (empty), had no infectivity (Fig. 2A).

In the production of pseudotypes bearing the CHIKV-E or VSV-G, the pseudotypes infect surrounding cells transfected with envelope protein expression vectors, further increasing the number of producing cells bearing the CHIKVΔE-NlucG or CHIKVΔCE-NlucG genome. On the other hand, in the production of CHIKVΔE-NlucG/CoV-2-S (or BA1-S) pseudotypes, the produced pseudotypes cannot spread the infection to surrounding cells because there is almost no ACE2 expression in 293T cells, and thus pseudotype-producing cells did not increase (Fig. 1B). This may be one of the reasons that the titers of CHIKVΔE-NlucG/CoV-2-S (or BA1-S) pseudotypes were lowest among the pseudotypes prepared in this study. Therefore, we tried to prepare the CHIKVΔE (or ΔCE)-NlucG/CoV-2-S (or BA1-S) pseudotype by using ACE2-expressing cells, but the cells transfected with the viral envelope expression vector, pCAGGS/CoV-2-S or pCAGGS/CoV-2-BA1-S, formed large fusion cells in poor condition, so the pseudotypes could not be generated (Supplemental figure 5).

Of course, in order to produce high-titer pseudotypes, it is crucial to produce a large number of cells expressing the vector RNA. Therefore, it is thought that pseudotype production can be efficiently increased by using transfection reagents that can efficiently introduce vector DNA into target cells or by using cell lines that show higher transfection efficiency. It is expected that future studies will continue to improve this experimental system.

Next, according to the manufacturer’s protocol, we tried to detect the infectivity of these pseudotypes by measuring Nluc activity using the Nano-Glo® Luciferase Assay System (Promega Corporation, Madison, WI). However, the background values were generally high and sometimes deviated from the number of GFP-positive cells detected (Fig. 2B-1). The deviation may have been due to the existence of free NLuc, free NLuc still fusing capsid protein or NLuc packaged in some kinds of particles that were released into the supernatant(Fig. 2B-2).

We therefore constructed firefly luciferase (luc)-expressing pseudotypes using the pCHIKVΔE-luc vector plasmid and viral envelope expression vector plasmids in the same manner as CHIKVΔE-NlucG/viral envelope protein pseudotypes (Supplemental figure 3). The backgrounds of firefly luciferase activity of the CHIKVΔE-luc/-inoculated cells were lower than that of Nluc, and the infectivities of these CHIKVΔE-luc/viral envelope protein pseudotypes were similar to the FFU of CHIKVΔE-NlucG/viral envelope protein pseudotypes (Fig. 2; Supplemental figure 3).

The CHIKVΔE-NlucG/VSV-G and CHIKVΔE-NlucG/CoV-2-S (or BA1-S) pseudotypes infectivities were neutralized with specific anti-envelope antibodies against each envelope protein (Fig.3), suggesting that the infections of CHIKV pseudotypes were strictly dependent on compatibility between the viral envelope protein and its receptor and there was no false-positive background infection.

**Fig. 3.**
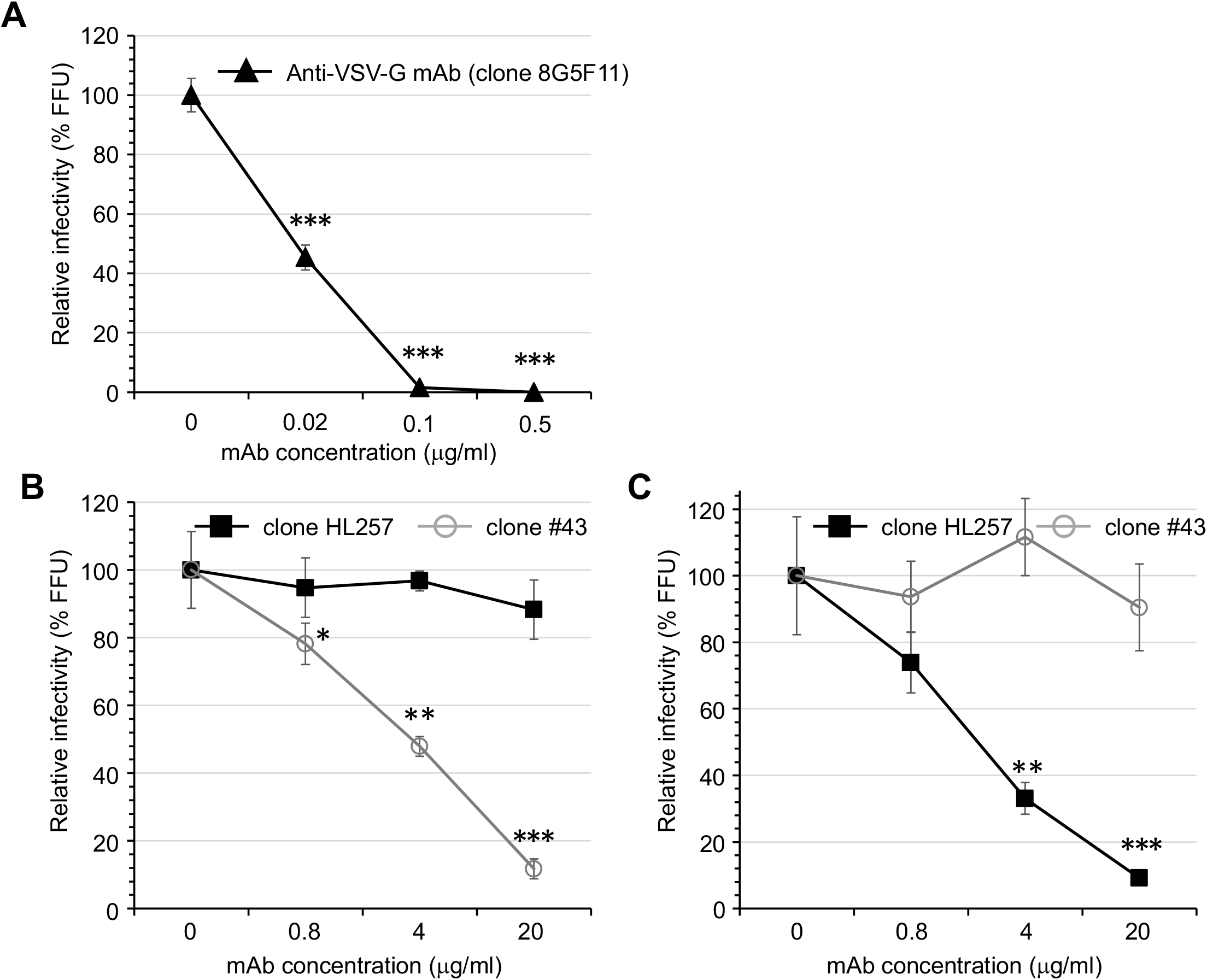
Neutralizing activity of anti-viral envelope protein detected by using the CHIKVΔE-NlucG/viral envelope pseudotype. Neutralizing activity of anti-VSV-Glycoprotein mAb (clone 8G5F11), and anti-SARS-CoV-2 spike protein mAbs (clone HL257 and clone #43) (see the Materials and Methods section) detected by neutralization assay using the (A) CHIKVΔE-NlucG/VSV-G, (B) CHIKVΔE-NlucG/CoV-2-S and (C) CHIKVΔE-NlucG/CoV-2-BA1-S, respectively. The plots shown are representative of three independent experiments performed in duplicate or triplicate. Data shown are expressed as the mean ± SD. Statistical significance was evaluated by an unpaired two-tailed *t* test. *Significant at *P* <0.05; **Significant at *P* <0.01; ***Significant at *P* <0.001.

In this study, we found that the anti-SARS-CoV-2 spike protein recombinant rabbit monoclonal antibody clone HL257 recognizes both the spike antigen of prototype CoV-2 and Omicron BA.1 (Supplemental figure 4) but neutralizes only CHIKVΔE-NlucG/CoV-2-BA1-S (Fig. 3). Therefore, SARS-CoV-2-BA.1 escaped neutralization of HL257 by changing the secondary or tertiary structure of its receptor binding domain (RBD).

In addition to BHK and BHK/hACE2 cells, pseudotyped infectivity was examined in Vero and Calu3 cells, which express ACE2 and are susceptible to SARS-CoV-2. Vero cells were more susceptible to infection by native-form pseudotype, CHIKVΔE-NlucG/CHIKV-E, than BHK and BHK/hACE2 cells, while the CHIKVΔE-NlucG/VSV-G infectivity was higher in BHK and BHK/hACE2 cells than Vero cells (Fig. 4). These results indicate that the higher susceptibility of BHK/hACE2 cells of CHIKVΔE-NlucG/CoV-2-S (or CoV-2-BA1-S) compared to Vero cells was not responsible for the capacity for CHIKV gene expression, but was responsible for the results of different infectivity via envelope proteins bearing pseudotypes (Fig. 4). BHK/hACE2 cells were more susceptible to infection of CHIKVΔE-NlucG/CoV-2-S than Vero cells that similar observed in native SARS-CoV-2 infection (Supplemental figure 2) also support these results. In addition, a difference in infectivity was observed between CHIKVΔE-NlucG/CoV-2-S and CHIKVΔE-NlucG/CoV-2-BA1-S in Vero cells but not in BHK/hACE2 or Calu3 cells, suggesting that the spike protein of SARS-CoV-2 Omicron BA.1 showed slightly higher binding affinity with African green monkey ACE2 than the spike protein of the prototype Wuhan strain SARS-CoV-2. It has been reported that the omicron RBD (strain BA.1) binds to ACE2 more strongly than does the prototypic RBD from the original Wuhan strain[55] and its binding affinity of hACE2 and RBD increased in the order of wild type (Wuhan-strain) < Beta < Alpha < Gamma < BA.1 [56, 57]. Recently, Tachibana et al. reported that the SARS-CoV-2 Omicron BA.1 showed a significantly lower binding affinity with human ACE2 than prototype SARS-CoV-2[58]. When pseudotype titers were adjusted for the titers in Vero cells, the relative infectivity of CHIKVΔE-NlucG/CoV-2-S to the human ACE2-expressing cells was higher than that of CHIKVΔE-NlucG/CoV-2-BA1-S, and the binding affinity of CoV-2-S to hACE2 appeared to be higher than that of CoV-2-BA1-S. Unfortunately, however, the absolute binding affinity of actual virus particles could not be determined in our system. Either way, the difference in binding affinity for hACE2 between CoV-2-S and CoV-2-BA1-S does not appear large enough to affect wild-type virus growth.

**Fig. 4.**
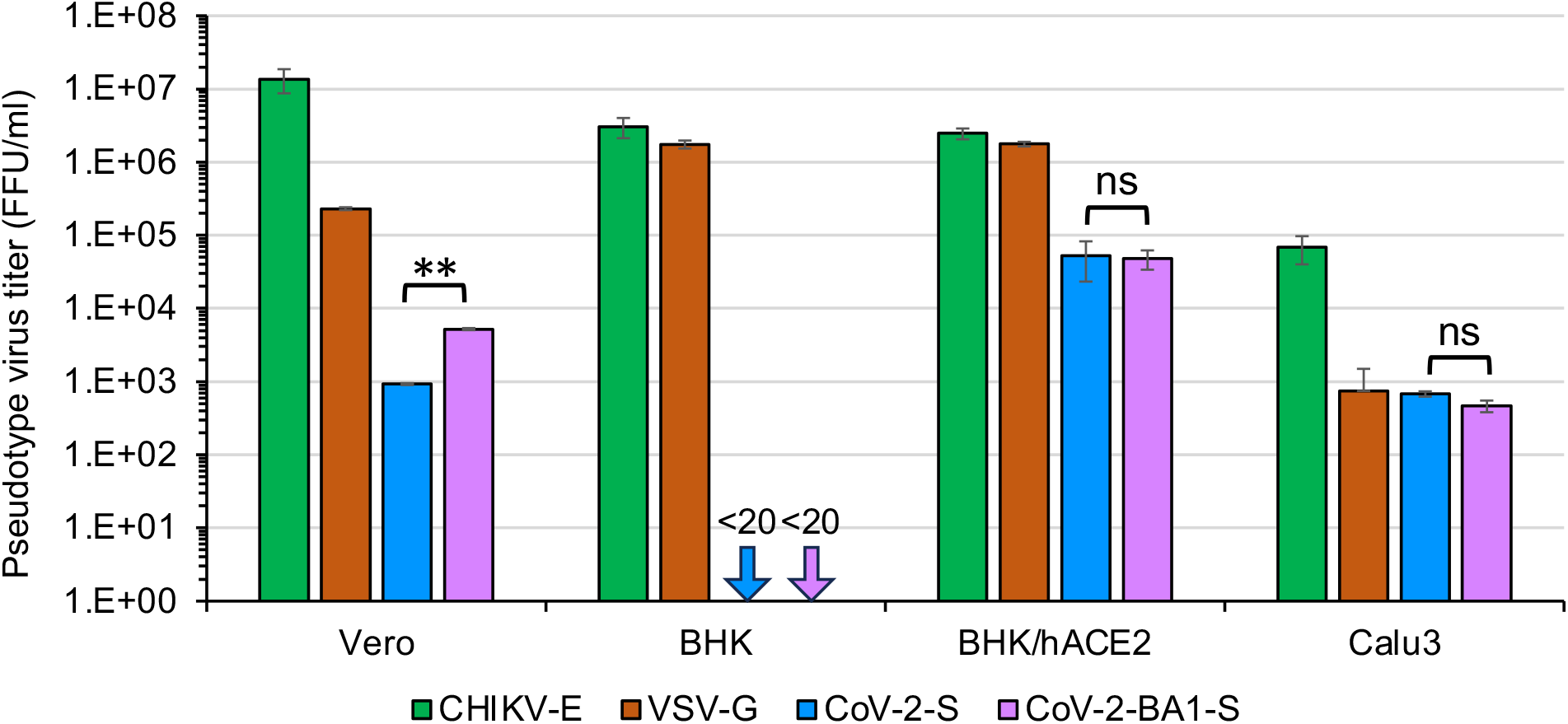
Susceptibilities of cultured cell lines to CHIKVΔE-NlucG/viral envelope protein pseudotypes. Vero, BHK, BHK-hACE2 and Calu-3 cells that had been plated 2 days earlier were inoculated with serially diluted pseudotype viruses and the titer of pseudotype viruses were determined by counting the GFP foci detected by fluorescence microscopy at 1 day after the inoculation. The pseudovirus titers (FFU/ml) shown are representative of three independent experiments performed in duplicate or triplicate and are expressed as the mean ± SD. Statistical significance was evaluated by an unpaired two-tailed *t* test. **Significant at *P* <0.01; ns: not significant.

Taken together, these results indicate that the infection of the pseudoviruses produced by our pseudotype virus system strictly depends on the expression of viral membrane proteins and viral receptors in target cells, there is no nonspecific infection, and the infectious titer is sufficient for virological studies. Our established pseudotype virus system can be used as a robust platform to study various virus infections and for screening and in-depth evaluation of neutralizing antibodies and antiviral agents.

Finally, it should be noted that the titer of the native-form CHIKV pseudotype, CHIKVΔCE-NlucG/CHIKV-E, was decreased by more than one-tenth compared to that of CHIKVΔE-NlucG/CHIKV-E, which was similar to the findings of previous studies [29, 30]. On the other hand, the titers of the pseudotype CHIKVΔCE-NlucG/VSV-G were reduced by only half compared to CHIKVΔE-NlucG/VSV-G, and the titers of CHIKVΔCE-NlucG/CoV-2-S were also decreased by only one-fifth compared to those of CHIKVΔE-NlucG/CoV-2-S (Fig.2-B-1). These results indicate that the titer of CHIKV-based pseudotypes bearing non-alphavirus envelope proteins is less dependent on the presence or absence of capsid proteins. One important implication of our findings is that alphavirus-derived replicon RNA vaccines, such as the constructs reported in Erasmus et al. [59], may have replication-competent viral activity. Therefore, we should exercise due caution in using replicon-type vaccine constructs expressing membrane proteins.

## ACKNOWLEDGMENTS

This work was supported by a JSPS KAKENHI grant (grant number 21K08490 to AT).

## Conflicts of interest

The authors declare that there are no conflicts of interest.

## Notes

### Competing Interest Statement

The authors have declared no competing interest.

