## supplemental figures for "Establishment of a Chikungunya virus pseudotype system strictly dependent on viral protein expression": Supplemental Figure .pdf

Supplemental figure 1

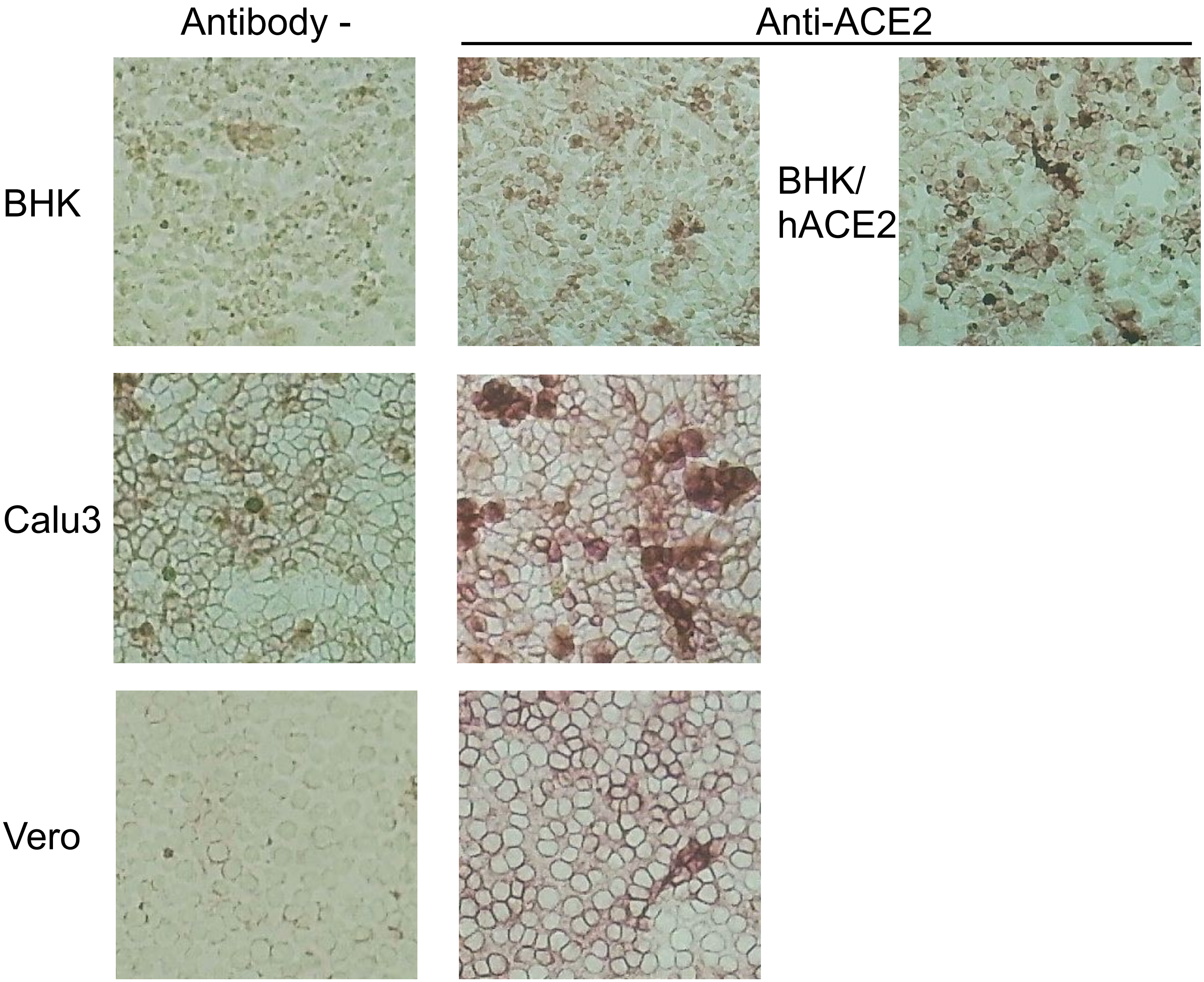

Sapplemental figure 2

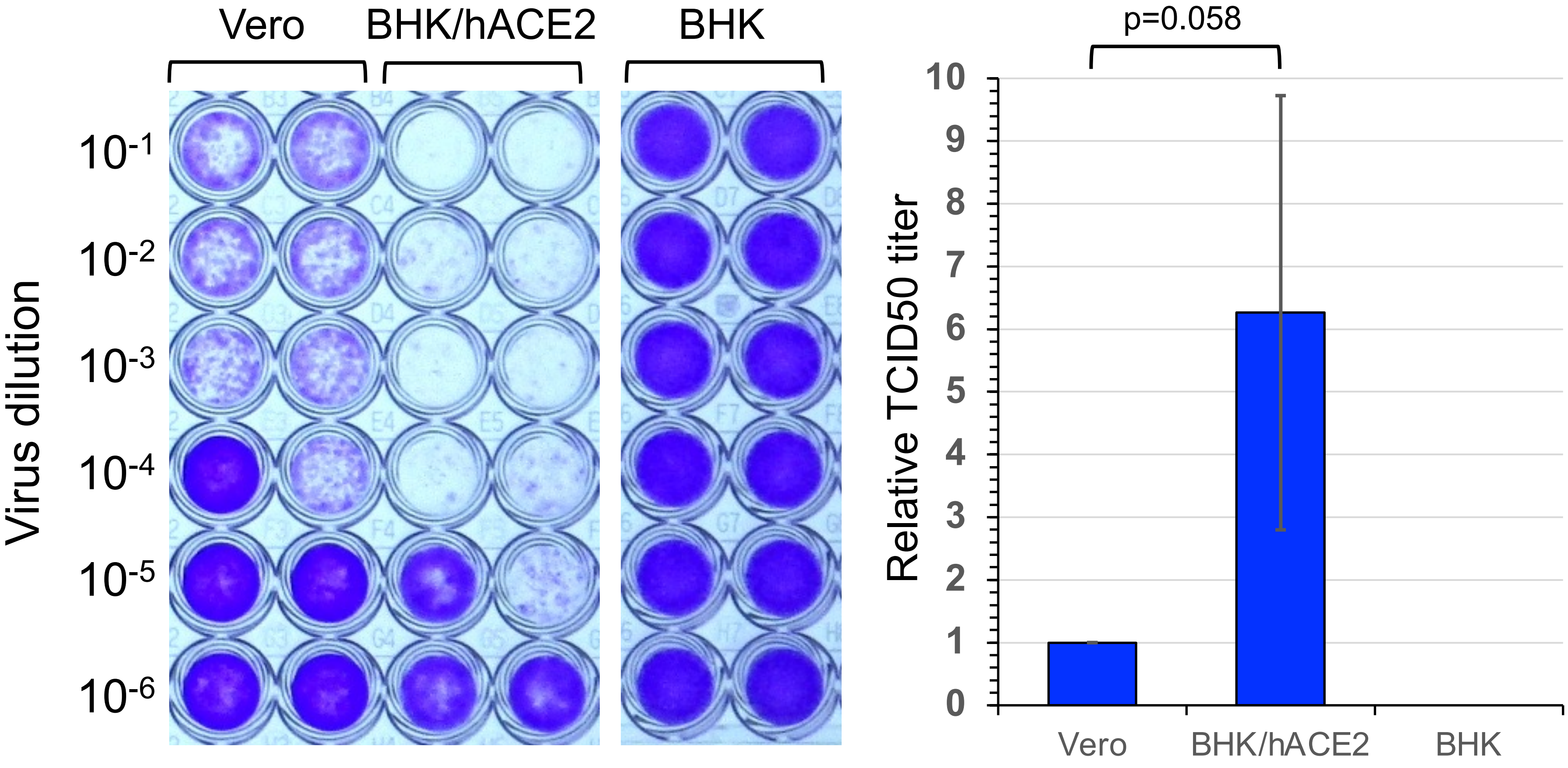

Sapplemental figure 3

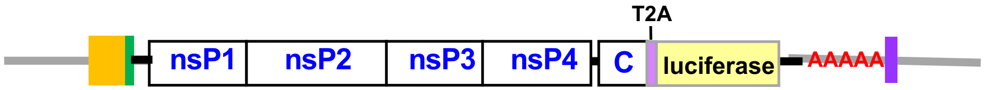


pCHIKVΔE-(Luc) expression vector plasmid

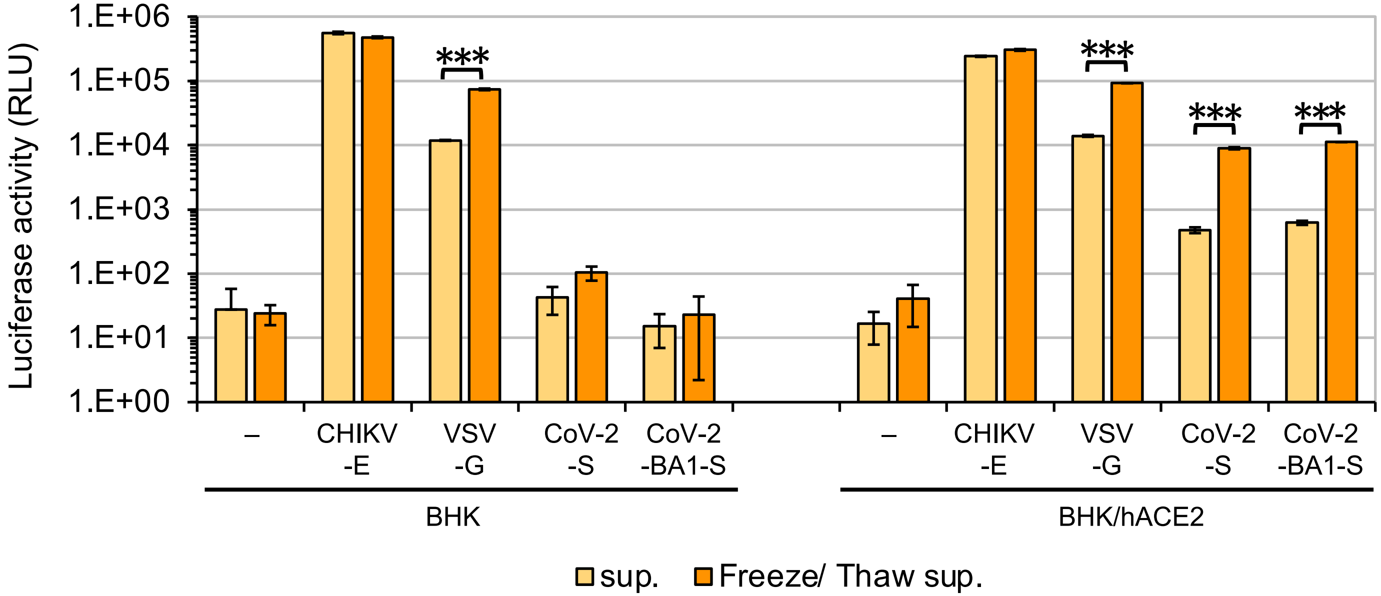

Sapplemental figure 4

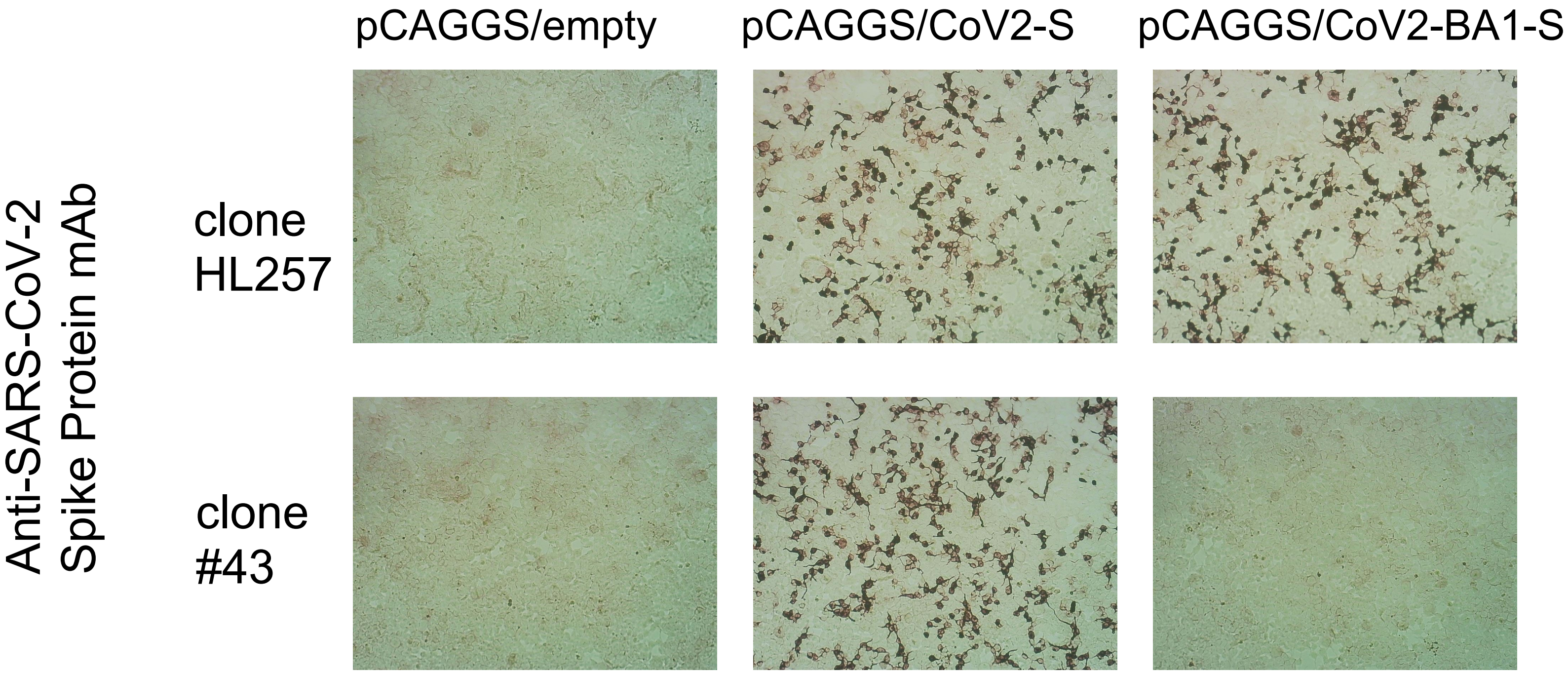

Sapplemental figure 5

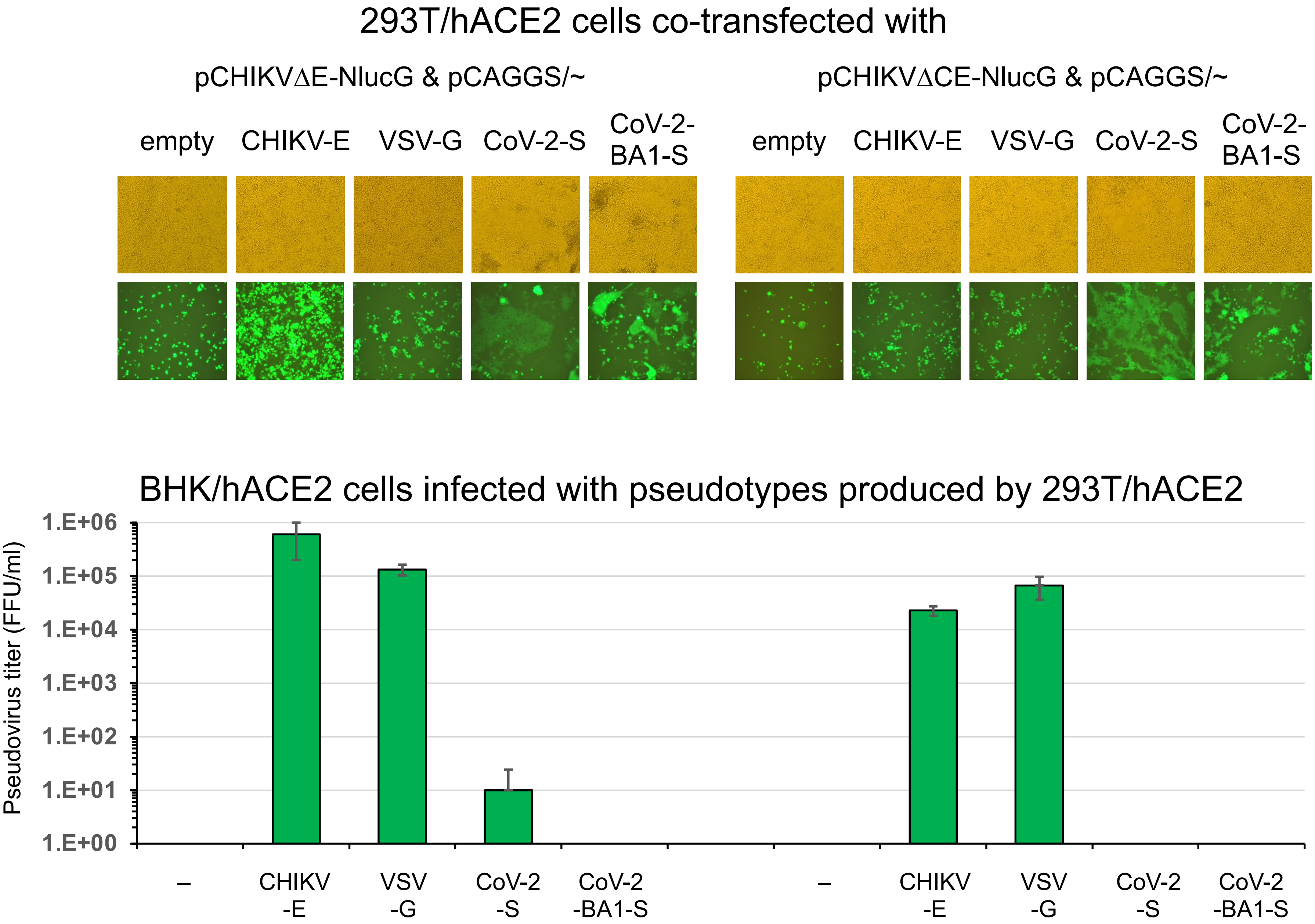

**Supplemental Figure 1**

Human ACE2 (hACE2) expressions in culture cells. BHK, BHK/hACE2, Calu3 and Vero cells were detected by using anti-ACE2 rabbit polyclonal antibody (catalogue no. 21115-1-AP; Proteintech Group) and the VECTASTAIN Elite ABC system (catalogue nos. PK-6100, SK-4600 and BA-1400).

**Supplemental Figure 2**

Vero, BHK and BHK/hACE2 cells were infected with serially diluted SARS-CoV-2 wuhan strain. The susceptibility of each cell line is shown by the virus titer (TCID50; determined using a Reed-Muench calculation) at 5 days after inoculation.

**Supplemental Figure 3**

Detection of luc activity expressed in BHK and BHK/hACE2 cells infected with CHIKVΔE-luc/(empty, CHIKV-E, VSV-G or CoV-2-S) pseudotypes using the Steady-Glo Luciferase Assay System (Promega Corporation, Madison, WI). The luc activities shown are representative of three independent experiments and are expressed as the mean ± SD. Statistical significance was evaluated by an unpaired two-tailed *t* test. ***Significant at $P$ <0.001.

**Supplemental Figure 4**

Reactivity of anti-SARS-CoV-2 spike protein mAbs (clone HL257 and #43) against the SARS-CoV-2 spike protein of the wuhan strain and BA.1 strain. 293T cells transfected with pCAGGS/empty, pCAGGS/CoV2-S or pCAGGS/CoV-2-BA1-S were reacted with anti-SARS-CoV-2 spike protein monoclonal antibodies, clone HL257 or clone #43 and detected by using the VECTASTAIN Elite ABC system (catalogue no. PK-6100, SK-4600 and BA-1400).

**Supplemental Figure 5**

**Few CHIKVΔE (or ΔCE)-NlucG/viral envelope protein pseudotypes produced by hACE2-expressing 293T cells**.

The hACE2-expressing 293T cells, 293T/hACE2, were co-transfected with pCHIKVΔE (or ΔCE)-NlucGFPbsr expression vector plasmid and pCAGGS/empty, pCAGGS/CHIKV-E, pCAGGS/VSV-G, pCAGGS/CoV-2-S or pCAGGS/CoV-2-BA1-S vector plasmid, and then the GFP expression in these transfected cells was detected by fluorescence microscopy at 3 days after transfection. The pseudotype samples were harvested as F/T sup as described in the Fig. 1 and then were inoculated into BHK-hACE2 cells, and GFP-expressing foci were detected by fluorescence microscopy at 1 day after inoculation as described in Fig. 2. Data shown are representative of three independent experiments and are expressed as the mean ± SD.
